# 3dSpAn: An interactive software for 3D segmentation and analysis of dendritic spines

**DOI:** 10.1101/864587

**Authors:** Nirmal Das, Ewa Baczynska, Monika Bijata, Blazej Ruszczycki, Andre Zeug, Dariusz Plewczynski, Punam Kumar Saha, Evgeni Ponimaskin, Jakub Wlodarczyk, Subhadip Basu

## Abstract

Three dimensional segmentation and analysis of dendritic spines involve two major challenges: 1) how to segment individual spines from the dendrites and 2) how to quantitatively assess the morphology of individual spines. We developed a software named 3dSpAn to address these two issues by implementing our previously published 3D multiscale opening algorithm in shared intensity space and using effective morphological features for individual dendritic spine plasticity analysis. 3dSpAn consists of four modules: Preprocessing and ROI selection, Intensity thresholding and seed selection, Multiscale segmentation and Quantitative morphological feature extraction. We show the results of segmentation and morphological analysis for different observation methods, including *in vitro* and *ex vivo* imaging with confocal microscopy, and *in vivo* samples, using high-resolution two-photon microscopy. The software is freely available, the source code, windows installer, the software manual and video tutorial can be obtained from: https://sites.google.com/view/3dSpAn/.

## 1. Introduction

In this article, we introduce an interactive software, 3dSpAn, designed to segment and quantitatively assess 3D morphology of dendritic spines. We evaluate the software performance with different imaging conditions (*in vitro, ex vivo* and *in vivo*). The software is based on an algorithm presented in our previous work [1], which provides a detailed theoretical background and validation of the proposed method with respect to the available state-of-the-art tools, such as Imaris software [2], demonstrating a high multi-user reproducibility. The developed software uses 3D multiscale opening (MSO) algorithm [3] to segment the spines from the dendritic segments.

The present paper is an application note, focusing on the software usage, and aiming to reduce user interaction in the segmentation process, and to help with estimation of adjustable segmentation parameters, with enhanced visualization. The software is freely available under GNU Lesser General Public License version 3 (LGPL V3). The software executable (3dSpAn V1.2 Installer for Windows), the source code, the detail user guide and video tutorials are available at: https://sites.google.com/view/3dSpAn/. A comprehensive user manual is also given as a supplementary document *3dSpAn_Supplementary* (also available as Manual in: https://sites.google.com/view/3dSpAn/). The complete workflow of the 3dSpAn software is described by the Figure S1-Figure S33 and Table S1 in the *3dSpAn_Supplementary*. Additionally, Video S1, Viedo S2 and Video S3 will describe how to perform segmentation and analysis of individual dendritic spine in 3dSpAn and how to visualize the segmented spines in 3D.

Dendritic spines are small membranous protrusions on neuronal dendrites having distinct structural features [4], controlling electrical and biochemical compartmentalization and playing major roles in activity and signal transmission of neural circuits [5]. The shape of dendritic spines changes spontaneously, or in response to neuronal stimulation [6,7]. These changes are related with learning and memory [8] and many neuropsychiatric and neurodegenerative diseases [9,10] e.g. Alzheimer’s disease [11], schizophrenia [12]. Many aspects of the existing structure-function relationship in dendritic spines are still unknown due to their complex morphology [8,10].

It is still challenging to segment individual spine and find exact spine boundaries, especially for lower-resolution microscopic images (when the possible highest resolution cannot be achieved, e.g. in *in vivo* imaging), that create difficulties in accurate modeling of 3D morphology of individual spine. The existing methods of segmentation and morphological analysis of dendritic spines either use 2D maximum intensity projection (MIP) image obtained from microscopic 3D image, or directly using microscopic 3D image. For example, the method presented in [13–15,21] use 2D MIP images for quantitative assessment of morphological changes in dendritic spines. However, 2D MIP images are misleading due to the loss of information and structure overlapping. Therefore, accurate quantitative morphological analysis of individual dendritic spine from 2-D MIP images is nearly impossible. Hence, the presented software segments individual dendritic spine directly in 3D and performs a morphological analysis of segmented dendritic spines. Several studies in the literature addressed the issue of segmentation and morphological analysis of dendritic spines directly for 3D microscopic images. A commercially available tool Imaris [2] allows user for 4D analysis of dendritic spine. Imaris is good for analyzing overall spine population but it fails to assess individual spine morphometry. In [17], Swanger et al proposed a method for automatic 4D analysis of dendritc spine morphology which follows the same pipeline as Imaris, however it does not quantify individual spine morphology. Several other methods are based on conventional machine learning as well as deep learning for segmentation and analysis of dendritic spine from microscopic images have been reported in the literature [32–34] but none of them is truely a 3D method. The main disadvantage of using deep learning based methods for segmentation and analysis of dendritic spines in 3D is the absence of sufficient manually annotated data in 3D. The works in [35,36] used deep learning for reconstruction of synapses from 3D microscopic images and identification of 3D morphological motifs but these are not suitable for segmentation and analysis of individual dendritic spine in 3D. Thus, a software allowing for interactive segmentation and morphological analysis of individual dendritic spine in 3D is still missing.

## 2. Results and Discussion

We show the results of analysis performed with images originating from three different laboratory techniques, *in vitro, ex vivo*, and *in vivo*. The details of these techniques and imaging modalities are described in Table 1. The morphological feature values (volume, length, head width and neck length) of the segmented spines from confocal microscopic image of *in vitro* neuronal culture (refer Figure 4) are shown in Table 2. Figure 5 and Figure 6 show the segmentation results in *ex vivo* sample and *in vivo* sample respectively. Table 3 and Table 4 show the morphological feature values of the segmented spines in Figure 5 and Figure 6. Figure 7 shows different segmented regions in a single 3D image. We observed that the performance of the software, in segmentation and analysis of dendritic spines, remained unchanged regardless of the laboratory technique used for imaging, and imaging modalities.

**Table 1.** Details of different imaging protocols.

| Methods | *In-vitro* | *Ex-vivo* | *In-vivo* |
| --- | --- | --- | --- |
| Model | primary /dissociated neuronal culture | brain slices | living animal |
| Animal | rat | mouse | mouse |
| Dendritic spine visualization method | transfection with EGFP plasmid under the control of synapsin-1 promoter at 14 day in vitro (DIV) of hippocampal neurons | DiI staining on fixed tissue | 2-photon imaging by cranial windows of transgenic mouse line (Thy1-GFP) expressing GFP under the thy1 promoter allowing visualization of the neurons in prefrontal cortex |
|  | GFP fluorescence | DiI fluorescence (red fluorescence channel) | GFP/YFP fluorescence |
| Protocols used in: [1,16] | **Primary hippocampal cultures** prepared from post-natal Day 0 Wistar rats. Hippocampi were dissected on ice in dissociation medium DM (81.8mM Na<sub>2</sub>SO<sub>4</sub>, 30mM K<sub>2</sub>SO<sub>4</sub>, 5.8mM MgCl<sub>2</sub>, 0.25mM CaCl<sub>2</sub>, 1mM HEPES pH 7.4, 20mM glucose; 1mM kynureic acid; 0.001% Phenol Red) and incubated twice for 15min at 37°C with 100 units of papain solution. Papain activity was stopped by applying a trypsin inhibitor dissolved in DM. The hippocampi were washed three times in plating medium [PM; Minimum Essential Media; 10% fetal bovine serum and 1% penicillin-streptomycin] and triturated to homogeneous suspension and centrifuged for 10min 1000xg resulting in cell pellet suspended in plating medium. Cells were plated at density 120000 cells per 18mm diameter coverslip coated with 1mg/ml poly-L-lysine (Sigma-Aldrich) and 2.5µg/ml laminin (Roche). After 1.5h plating medium was exchanged for maintenance medium (MM; Neurobasal-A, 2% B-27 supplement and 1% penicillin-streptomycin). The cells were maintained at 37°C under a humidified 5% CO<sub>2</sub> atmosphere. | **Biolistic staining with DiI dye** was performed on C57BL/6J mouse brain slices. The mice were anesthetized and transcardially perfused with 1.5% paraformaldehyde. The brains were sliced for 140µm thick and allowed for 1.5h at room temperature. Random dendrite labeling was performed using 1.6mm tungsten particles (Bio-Rad) coated with propelled lipophilic fluorescent dye (1,1'-dioctadecyl-3,3,3',3'-tetramethylindocarbocyanine perchlorate, Invitrogen) delivered to the cells by gene gun (Bio-Rad). | **Virus injections** Implant cranial windows and inject HAPLN1-Scarlet + Synaptophysin-BFP AAVs in PFC for Thy1-GFP mice. Imaging after 4 weeks postinjection: Anaesthetize with Isoflurane to perform baseline images (3 to 4 positions) and 24hours after performance of single image for all positions. |
| Imaging protocols | **Live cell imaging** was performed at 20 – 22 DIV. Cultures were transfected at 14 DIV with Syn-GFP plasmid under a synapsin-1 promoter with Lipofectamine reagent. For imaging the cultures were placed into the acquisition chamber under 37°C temperature and 5% CO<sub>2</sub> conditions. The images of dendritic segments were acquired using Zeiss LSM780 confocal microscope with a PL Apo 40x/1.2 NA water immersion objective using 488nm diode-pumped solid-state laser at 10% transmission at a pixel count 1024x1024. Z-stacks were collected for each cell with a 0.4µm step size. An additional digital zoom was applied, resulting in a lateral resolution of 0.07µm. Transgenic mouse line (Thy1-GFP) expressing GFP under the thy1 promoter. | **Imaging:** Images of dendrites in different brain regions were acquired under 561nm fluorescent illumination using a confocal microscope (63x objective, 1.4 NA) at a pixel resolution of 1024 × 1024 with a 3.43 zoom, resulting in a 0.07µm pixel size [25]. | **Imaging:** Images of dendrites in different brain regions were acquired under 488nm fluorescent illumination using a 2-photon microscope (40x objective) with a 2.0 zoom. |
| Procedure | See [25,26] | See [25,27] | See [28–31] |
| Aparatus | Fluorescent confocal microscopy equipped with live cell imaging setup (Zeiss LSM780) | Fluorescent confocal microscopy (Zeiss LSM800) | 2-photon microscopy (Zeiss LSM 7 MP Axio Examiner.Z1) |

**Table 2.**
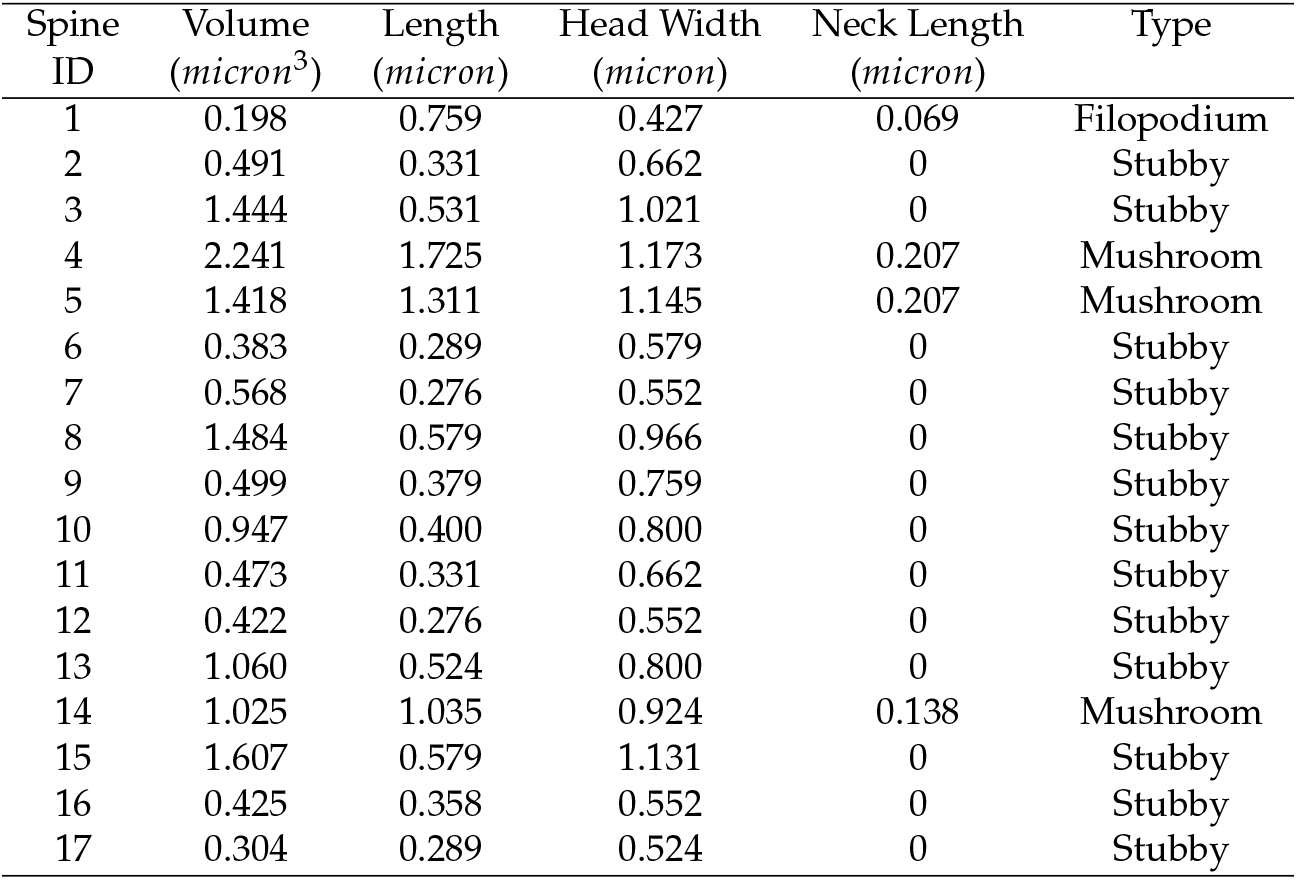
Morphological features (volume, length, head width, neck length) and types of dendritic spines for each spine shown in Figure 4(a).

| Spine ID | Volume ($\mu\text{m}^3$) | Length ($\mu\text{m}$) | Head Width ($\mu\text{m}$) | Neck Length ($\mu\text{m}$) | Type |
| --- | --- | --- | --- | --- | --- |
| 1 | 0.198 | 0.759 | 0.427 | 0.069 | Filopodium |
| 2 | 0.491 | 0.331 | 0.662 | 0 | Stubby |
| 3 | 1.444 | 0.531 | 1.021 | 0 | Stubby |
| 4 | 2.241 | 1.725 | 1.173 | 0.207 | Mushroom |
| 5 | 1.418 | 1.311 | 1.145 | 0.207 | Mushroom |
| 6 | 0.383 | 0.289 | 0.579 | 0 | Stubby |
| 7 | 0.568 | 0.276 | 0.552 | 0 | Stubby |
| 8 | 1.484 | 0.579 | 0.966 | 0 | Stubby |
| 9 | 0.499 | 0.379 | 0.759 | 0 | Stubby |
| 10 | 0.947 | 0.400 | 0.800 | 0 | Stubby |
| 11 | 0.473 | 0.331 | 0.662 | 0 | Stubby |
| 12 | 0.422 | 0.276 | 0.552 | 0 | Stubby |
| 13 | 1.060 | 0.524 | 0.800 | 0 | Stubby |
| 14 | 1.025 | 1.035 | 0.924 | 0.138 | Mushroom |
| 15 | 1.607 | 0.579 | 1.131 | 0 | Stubby |
| 16 | 0.425 | 0.358 | 0.552 | 0 | Stubby |
| 17 | 0.304 | 0.289 | 0.524 | 0 | Stubby |

**Table 3.**
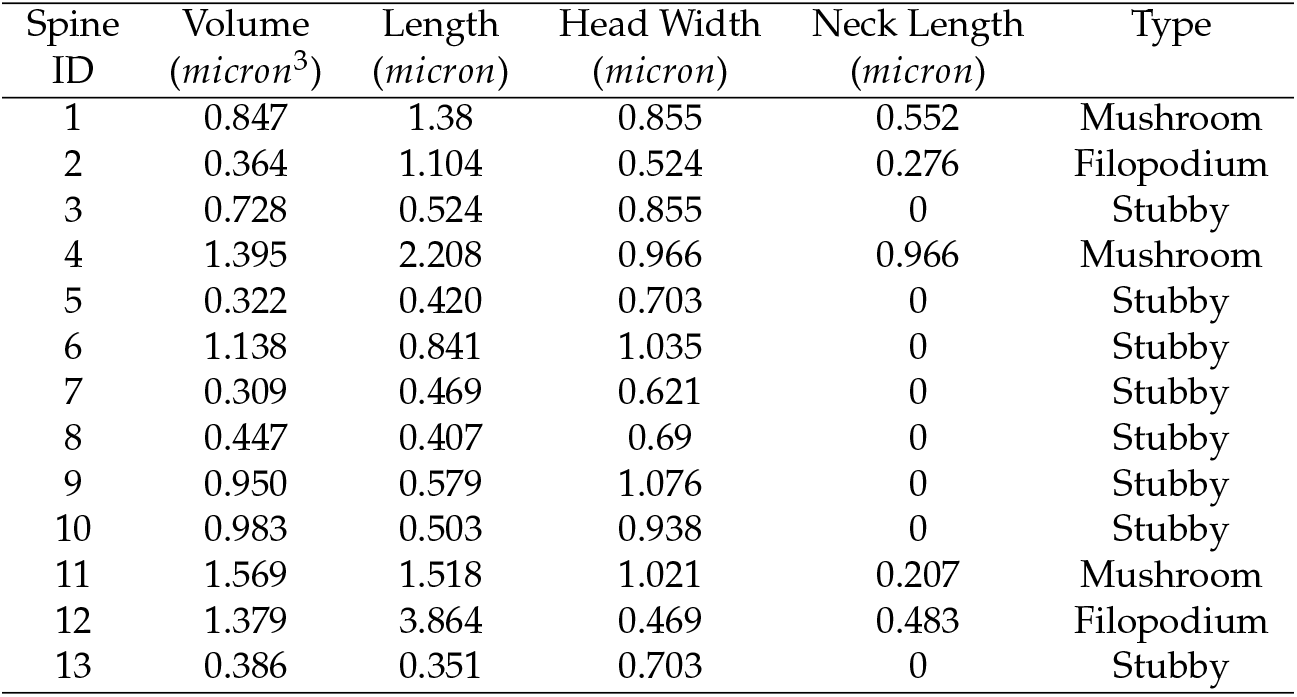
Morphological features (volume, length, head width, neck length) and types of dendritic spines for each spine shown in Figure 5

| Spine ID | Volume ($\mu\text{m}^3$) | Length ($\mu\text{m}$) | Head Width ($\mu\text{m}$) | Neck Length ($\mu\text{m}$) | Type |
| --- | --- | --- | --- | --- | --- |
| 1 | 0.847 | 1.38 | 0.855 | 0.552 | Mushroom |
| 2 | 0.364 | 1.104 | 0.524 | 0.276 | Filopodium |
| 3 | 0.728 | 0.524 | 0.855 | 0 | Stubby |
| 4 | 1.395 | 2.208 | 0.966 | 0.966 | Mushroom |
| 5 | 0.322 | 0.420 | 0.703 | 0 | Stubby |
| 6 | 1.138 | 0.841 | 1.035 | 0 | Stubby |
| 7 | 0.309 | 0.469 | 0.621 | 0 | Stubby |
| 8 | 0.447 | 0.407 | 0.69 | 0 | Stubby |
| 9 | 0.950 | 0.579 | 1.076 | 0 | Stubby |
| 10 | 0.983 | 0.503 | 0.938 | 0 | Stubby |
| 11 | 1.569 | 1.518 | 1.021 | 0.207 | Mushroom |
| 12 | 1.379 | 3.864 | 0.469 | 0.483 | Filopodium |
| 13 | 0.386 | 0.351 | 0.703 | 0 | Stubby |

**Table 4.**
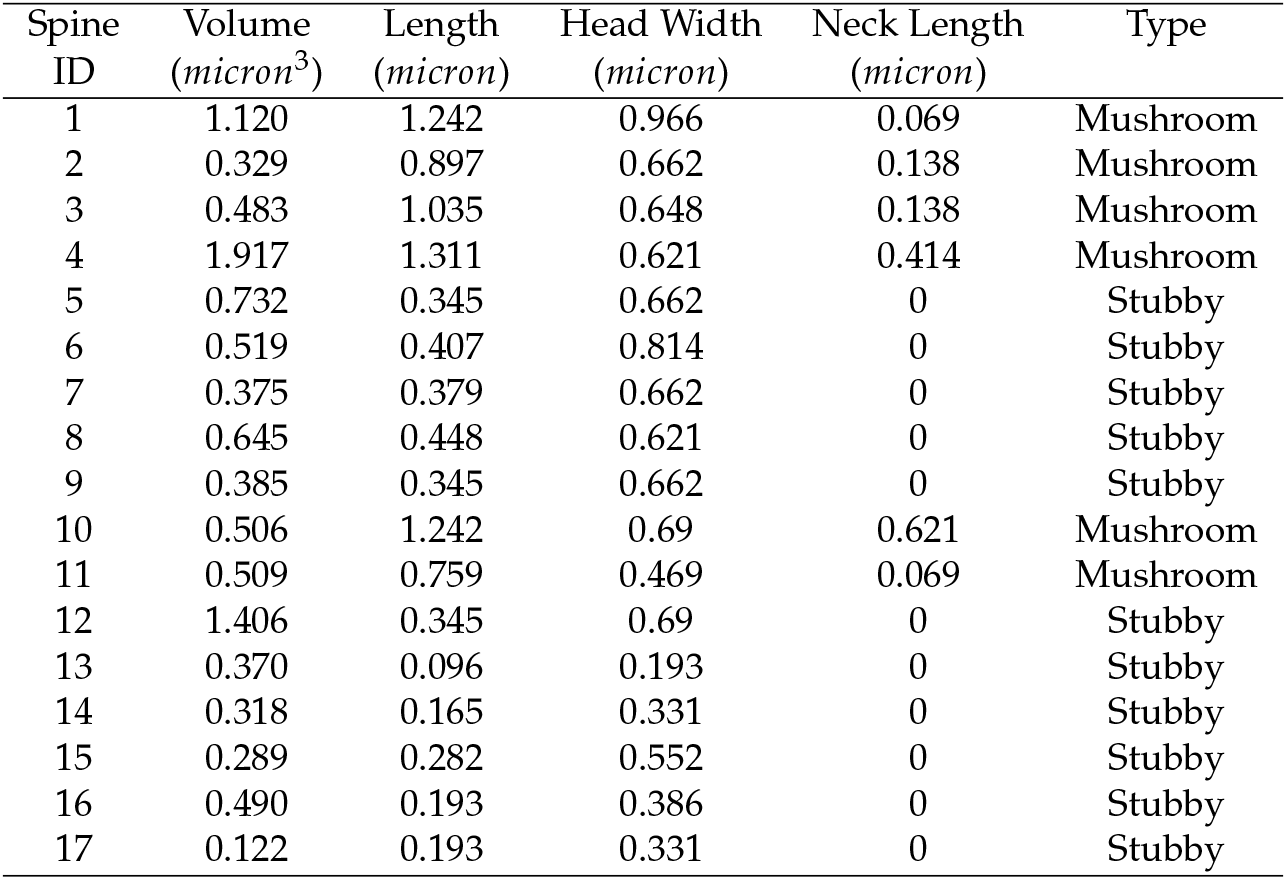
Morphological features (volume, length, head width, neck length) and types of dendritic spines for each spine shown in Figure 6.

| Spine ID | Volume ($\mu\text{m}^3$) | Length ($\mu\text{m}$) | Head Width ($\mu\text{m}$) | Neck Length ($\mu\text{m}$) | Type |
| --- | --- | --- | --- | --- | --- |
| 1 | 1.120 | 1.242 | 0.966 | 0.069 | Mushroom |
| 2 | 0.329 | 0.897 | 0.662 | 0.138 | Mushroom |
| 3 | 0.483 | 1.035 | 0.648 | 0.138 | Mushroom |
| 4 | 1.917 | 1.311 | 0.621 | 0.414 | Mushroom |
| 5 | 0.732 | 0.345 | 0.662 | 0 | Stubby |
| 6 | 0.519 | 0.407 | 0.814 | 0 | Stubby |
| 7 | 0.375 | 0.379 | 0.662 | 0 | Stubby |
| 8 | 0.645 | 0.448 | 0.621 | 0 | Stubby |
| 9 | 0.385 | 0.345 | 0.662 | 0 | Stubby |
| 10 | 0.506 | 1.242 | 0.69 | 0.621 | Mushroom |
| 11 | 0.509 | 0.759 | 0.469 | 0.069 | Mushroom |
| 12 | 1.406 | 0.345 | 0.69 | 0 | Stubby |
| 13 | 0.370 | 0.096 | 0.193 | 0 | Stubby |
| 14 | 0.318 | 0.165 | 0.331 | 0 | Stubby |
| 15 | 0.289 | 0.282 | 0.552 | 0 | Stubby |
| 16 | 0.490 | 0.193 | 0.386 | 0 | Stubby |
| 17 | 0.122 | 0.193 | 0.331 | 0 | Stubby |

## 3. Materials and Methods

3dSpAn comprises four main modules: Preprocessing and ROI selection (M1), Intensity thresholding and seed selection (M2), Multiscale segmentation (M3) and Quantitative morphological feature extraction (M4). Figure 1 describes an overall workflow of 3dSpAn, the different components of GUI and a snapshot of selected regions of interest (ROI), for a confocal microscopic image of *in vitro* neuronal culture is used. 3dSpAn software is implemented in C++ language and Qt development environment [20]. For 3D visualization of the segmented spines, we used the open source software ITK SNAP [19]

**Figure 1.**
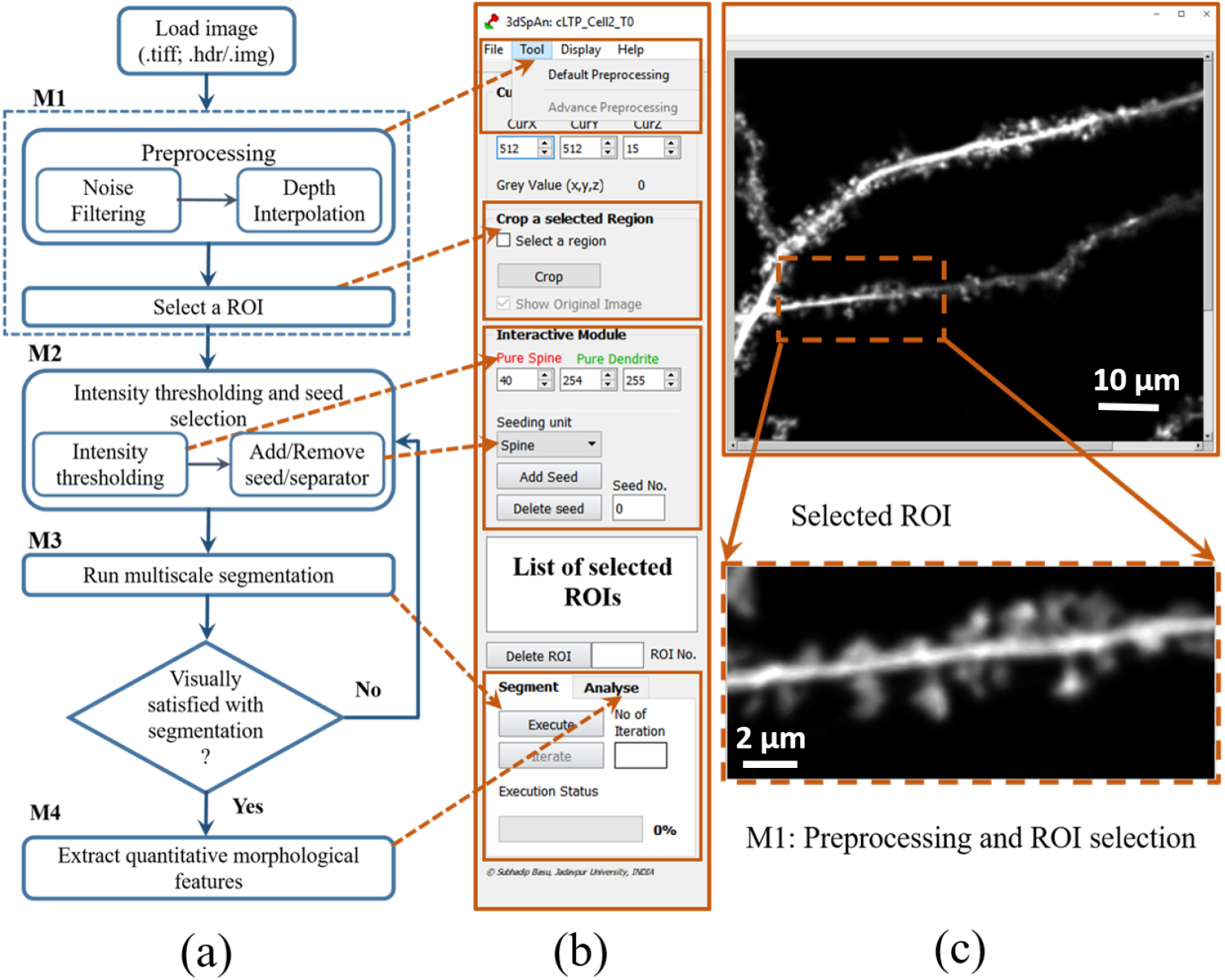
3dSpAn workflow and brief description of the GUI and the M1 module. (a) Diagram of 3dSpAn workflow. The blocks are representing four main modules: M1, M2, M3 and M4 (b) GUI working panel. The main modules are enclosed with rectangular boxes, dashed arrows show the correspondence with the appropriate blocks of the workflow diagram. (c) An image of *in vitro* neuronal culture is loaded in GUI (above) and a ROI is selected (dotted rectangular box). The magnified version of selected ROI (Scale bar=2 *μ*m) is shown below the full image.

### 3.1. Preprocessing and ROI selection (M1)

The preprocessing step is used to eliminate image noise (mainly salt and pepper noise) and to equalize image resolution along all three axes (confocal images have lower axial resolution than lateral resolution). 3D median filter [22] is applied to eliminate image noise (the median filter kernel size and voxel dimensions can be selected by the user, details are given in *3dSpAn_Supplementary*). Bilinear interpolation [23] is applied along the axial direction for appropriate scaling, producing smooth interpolating results in a real time. The user selects a region of interest (ROI) and for segmentation and quantitative morphological analysis of individual dendritic spine. The benefits of working with a smaller ROI is that the intensity thresholds are better estimated, and adjustment of these thresholds is also easier for a smaller ROI. After working on the current ROI, the user can select another ROI from the image and perform further segmentation and analysis. It is possible to stop in between the analysis process and to resume it later from the saved profile.

### 3.2. Intensity thresholding and seed selection (M2)

In order to perform the segmentation of dendritic spines, first we segment dendrite and spines together from the background, and then we segment individual spines from the dendrite. Generally, spines are of low intensity and dendrites are of higher intensity but spine and dendrite share a common intensity range. Two intensity thresholds for spine and dendrite, *th*_*s*_ and *th*_*d*_ respectively are initially estimated for the selected ROI (*R*_*i*_) in the following way. Let *μ* be mean intensity of the ROI and *δ* be standard deviation. The thresholds *th*_*s*_ and *th*_*d*_ are calculated as

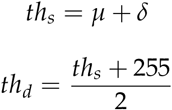

If intensity at some point *P*(*x, y, z*), *I*(*P*) < *th*_*s*_ then it is classified as a pure background point and if *I*(*P*) ≥ *th*_*d*_ then it is classified as a pure dendrite.The intensity range between *th*_*s*_ and *th*_*d*_ is the shared intensity space between spine and dendrite. A monotonically increasing fuzzy membership function is used here to calculate spine and dendrite membership (*μ*_*s*_ and *μ*_*d*_) of each pixel,

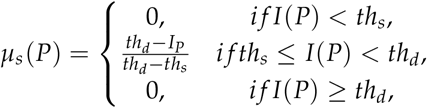

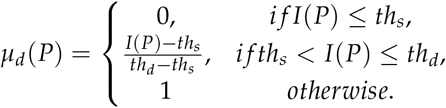

It may happen a spine in *R*_*i*_ are of high intensity (> *th*_*d*_) or a spine is disconnected from dendrite because of low intensity (< *th*_*s*_). In this case we have to modify the value of *th*_*s*_ and *th*_*d*_ to segment the spines, accordingly. For some ROIs, we may encounter spines with higher intensities, and almost equal to the intensity of dendrite. In this case we have to increase the value of *th*_*d*_. The intensity space between *th*_*s*_ and *th*_*d*_ can be visualized as color transition from red to green, changing from spine to dendrite. This color coded visualization helps the user to modify *th*_*s*_ and *th*_*d*_ manually. To select a dendritic spine (*S*_*i*_) for segmentation, the user needs to place a seed point on it. Multiple seed points can be placed for a single spine (*S*_*i*_). The pixels with intensities greater than *th*_*d*_ are considered as implicit dendritic seeds. Explicit dendritic seeds are required if *μ*_*d*_ values are too low for voxels belonging to dendrite region. In this case, a separator is placed to separate two touching spines. Figure 2 shows intensity thresholding and a seed selection module on the image presented in Figure 1.

**Figure 2.**
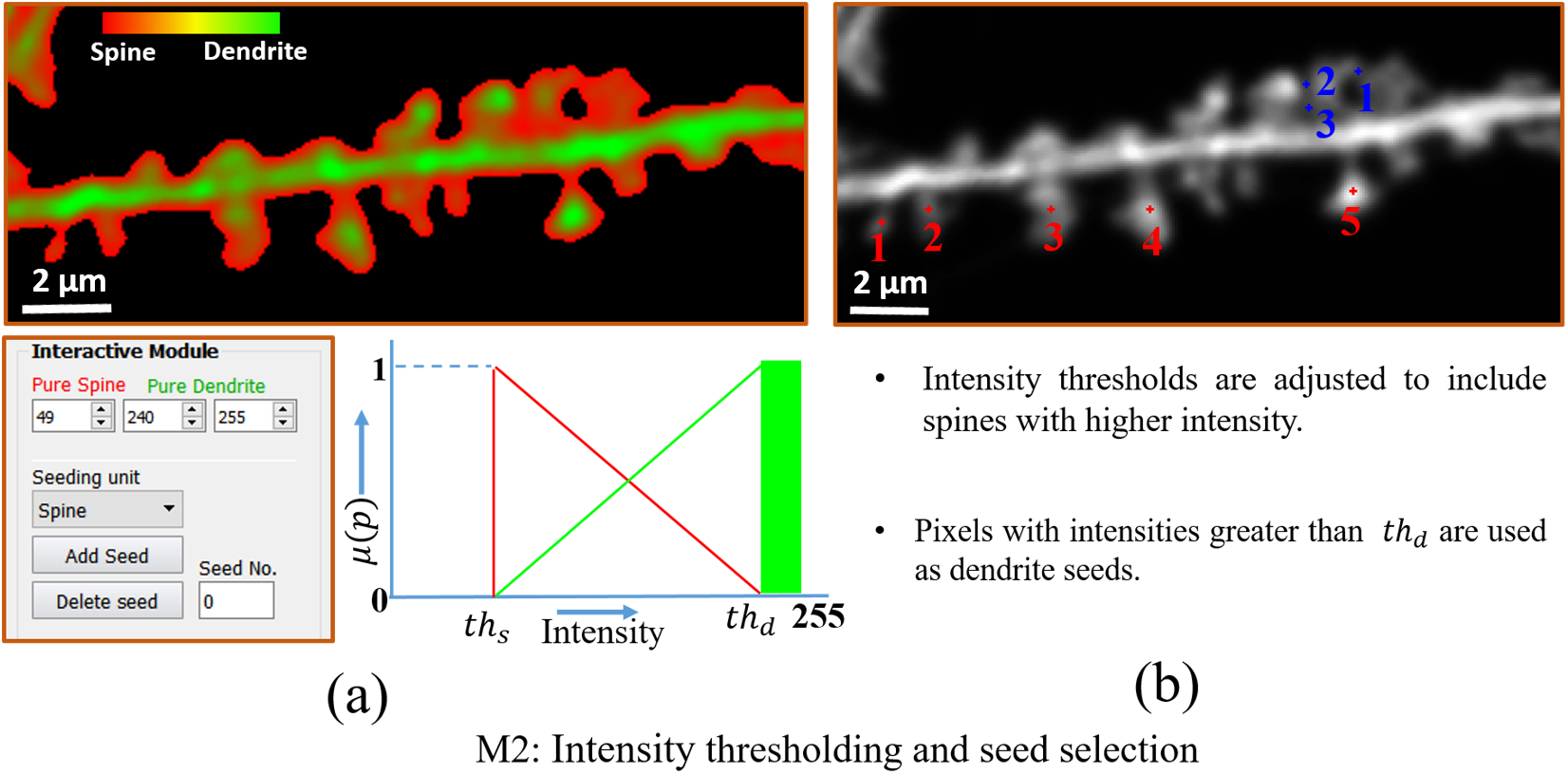
M2 module. (a) Visualization of the shared intensity space between spine and dendrite in the selected ROI (Scale bar=2 *μ*m) of Figure 1(c) with a color transition from red to green, describing the spine association and the dendrite association for the pixel intensity value (top). The block interactive module shows the intensity thresholds *th_s_*(40) and *th_d_*(240), for the spine and the dendrite respectively. The graph shows fuzzy membership curve for both spines (red) and the dendrite (green). (b) User-specified spine seeds (red) and separators (blue), placed at the same depth on different spines (Scale bar=2 *μ*m). The seeds are numbered in the order in which they were placed. Often, to include a spine, the user needs to adjust *th_s_* and *th_d_* values. The pixels with intensity value greater than *th_d_* are used as dendrite seeds implicitly. For this ROI, explicit dendrite seeds are not given.

### 3.3. Multiscale segmentation (M3)

The user-specified seed points and separators are considered as inputs for the MSO algorithm to segment the dendrites and the spines. MSO algorithm segments two conjoint objects, namely spines from dendrites in this case, coupled at unknown locations and at arbitrary scales in the shared intensity space (bounded by *th*_*s*_ and *th*_*d*_). With user-specified seed points and separators, the MSO algorithm separates the spines from the dendrites at a specific scale based on fuzzy distance transform (FDT) and fuzzy morphoconnectivity strength. After segmentation at the specific scale, the previous separation boundary is frozen using constrained morphological dilation, enabling segmentation at the next, finer scale. In this iterative approach of MSO, it takes several iterations to grow path-continuity of an object starting from its seed, often falling in large-scale regions, to a peripheral location with fine scale details, see Saha et. al [3] for theoretical and mathematical details of MSO algorithm. Figure 3 shows the GUI multiscale segmentation module.

**Figure 3.**
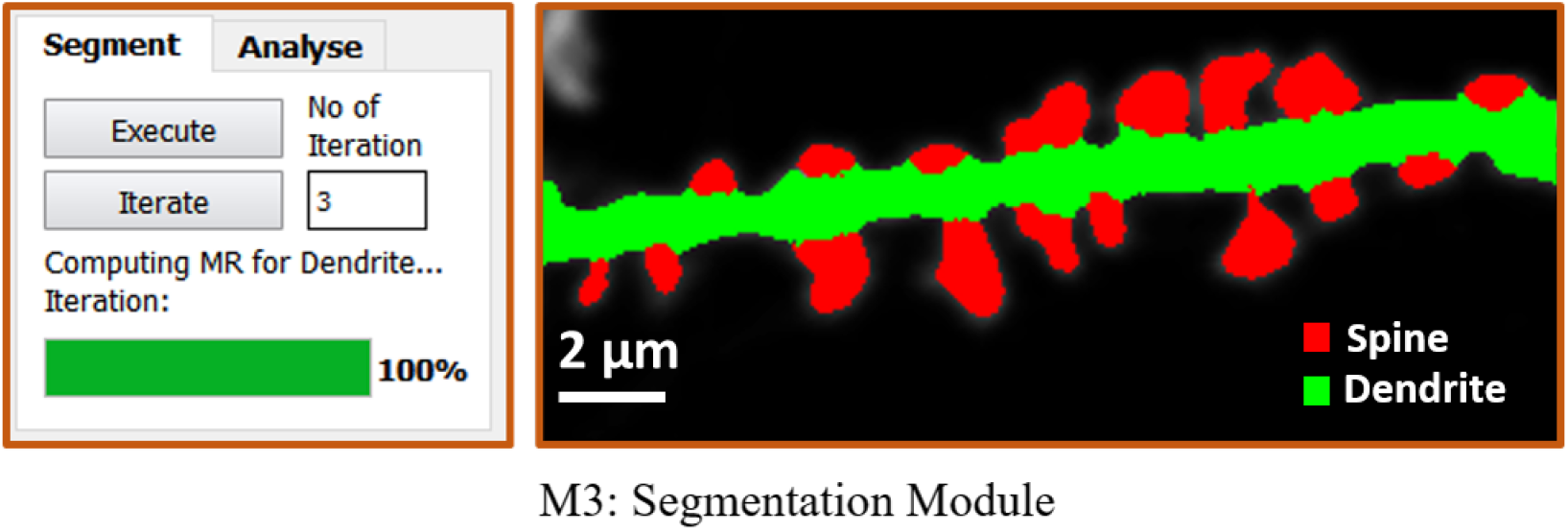
M3 module and segmented spines (Scale bar=2 *μ*m) from the selected ROI of Figure 1(c). The segmentation module of the GUI shows the number of times the MSO algorithm iterates to achieve the segmentation result.

### 3.4. Quantitative morphological feature extraction (M4)

After 3D segmentation, the feature extraction module extracts key morphological features for each of the segmented spine *S_i_*, like volume, length, head width, and neck length. These features are extracted by identifying three characteristic points for spines, 1) the central base point, i.e., the central point of the junction between the segmented spine and dendrite, 2) the central head point, i.e., the locally deepest point in the spine, and 3) the farthest point on the spine from the central base point, determined using FDT based shortest path. Using these features, each *S_i_* is categorized into one of the three major spine classes: stubby, filopodia or mushroom spines. For the mathematical detail of the quantitative measurements and spine classification please refer to our previous paper [16]. Figure 4 shows the quantitative morphological feature extraction module.

**Figure 4.**
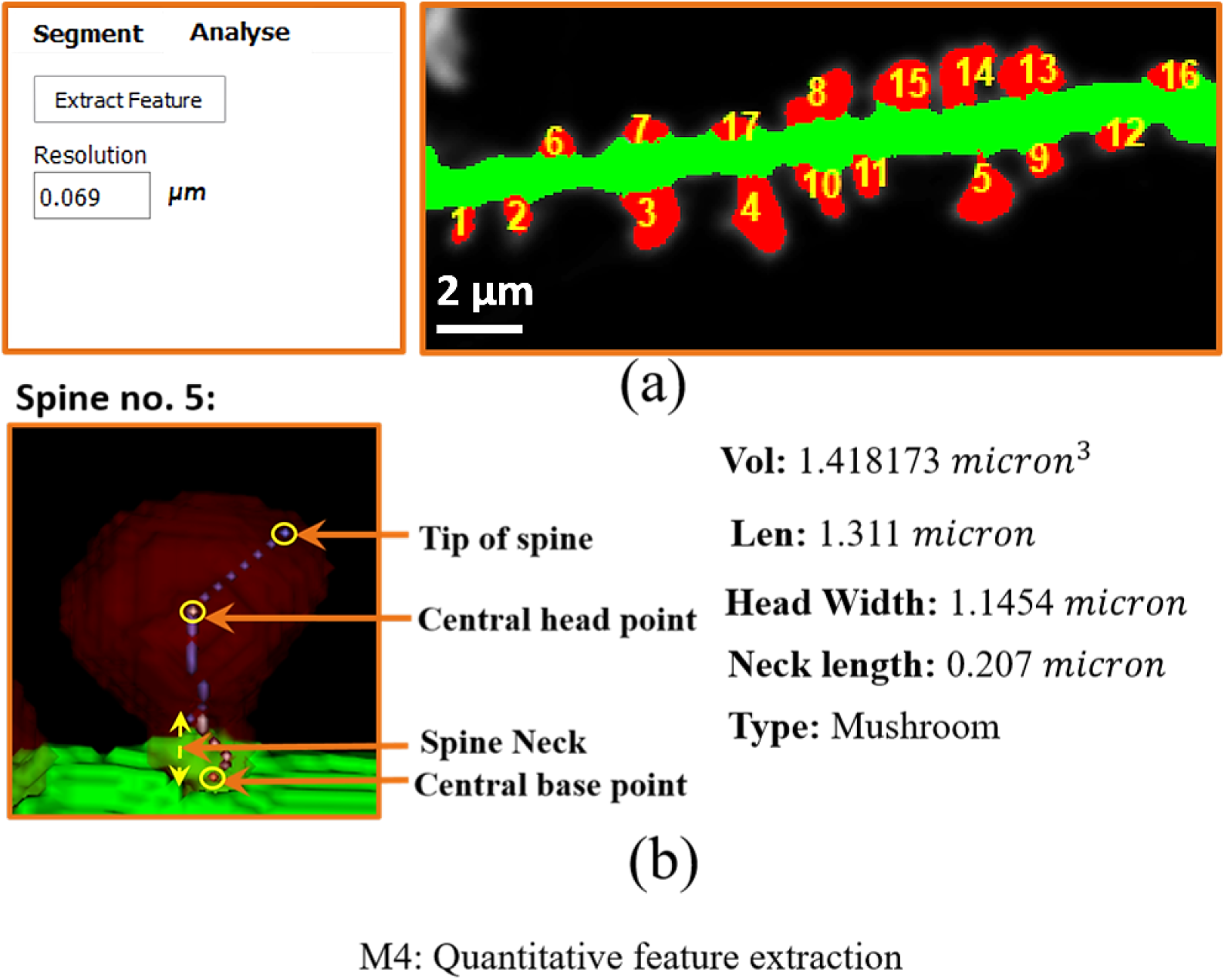
M4 module. (a) After quantitative morphological feature extraction, the segmented spines (Scale bar=2 *μ*m) (see Figure 3) are numbered in the same order as user-specified seed points. (b) Magnified and rotated 3D view of a spine no. 5. The spine is visualized with popular ITK-Snap software [19]. The transparent visualization helps to see the calculated path and points inside spine. The key points like *Central base point*, *Central head point* and *Tip of the spine* are marked (small yellow circle). Spine volume, spine length, head width and neck length are also calculated. Based on these measurements, spine classification is performed. Here the segmented spine belongs to the Mushroom class.

**Figure 5.**
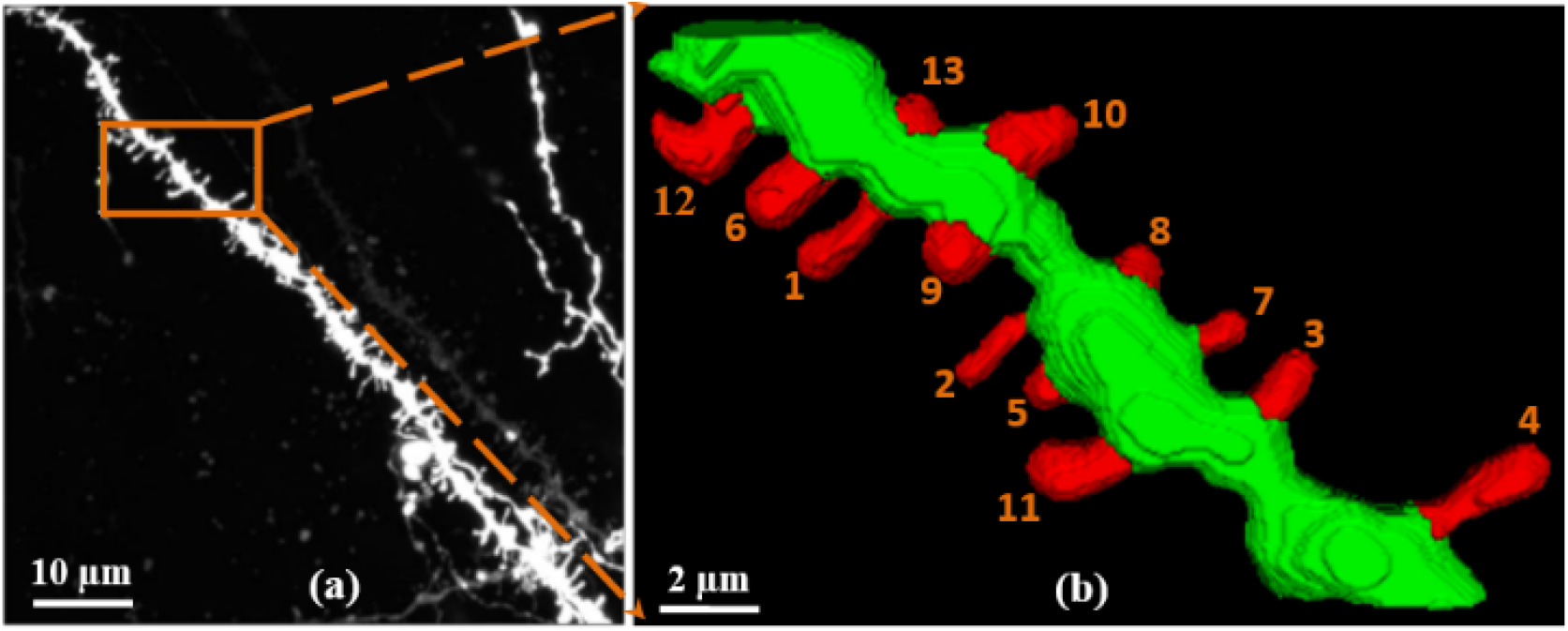
3D reconstruction of the segmented spines (*ex vivo*). (a) A ROI is selected (shown in rectangular box) in an *ex vivo* image (Scale bar=10 *μ*m) of mouse brain slices. (b) Magnification of 3D reconstruction of the segmented spine and dendrite with individual spine numbering in the selected ROI(Scale bar=2 *μ*m).

**Figure 6.**
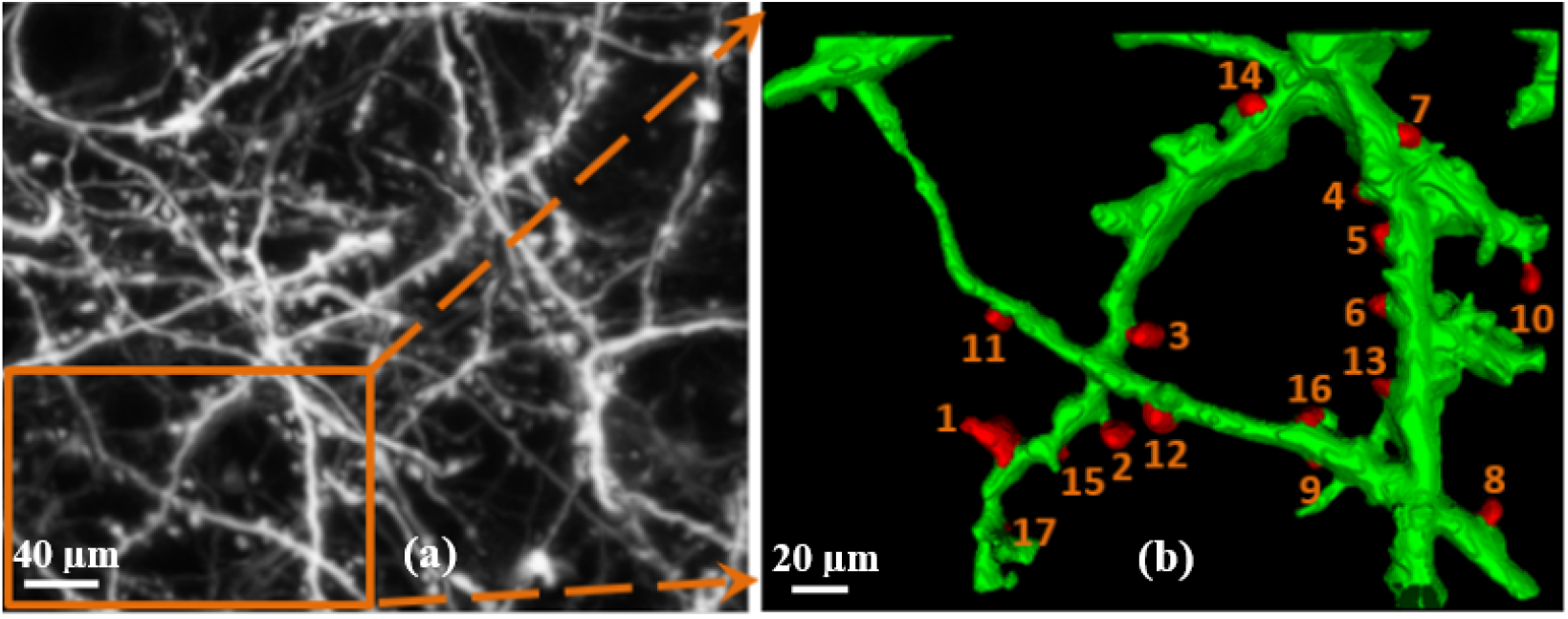
3D reconstruction of the segmented spines (*in vivo*). (a) A ROI is selected in an *in vivo* image (Scale bar=40 *μ*m) of living mouse brain. (b) Magnification of 3D reconstruction of the segmented spine and dendrite with individual spine numbering in the selected ROI (Scale bar=20 *μ*m).

**Figure 7.**
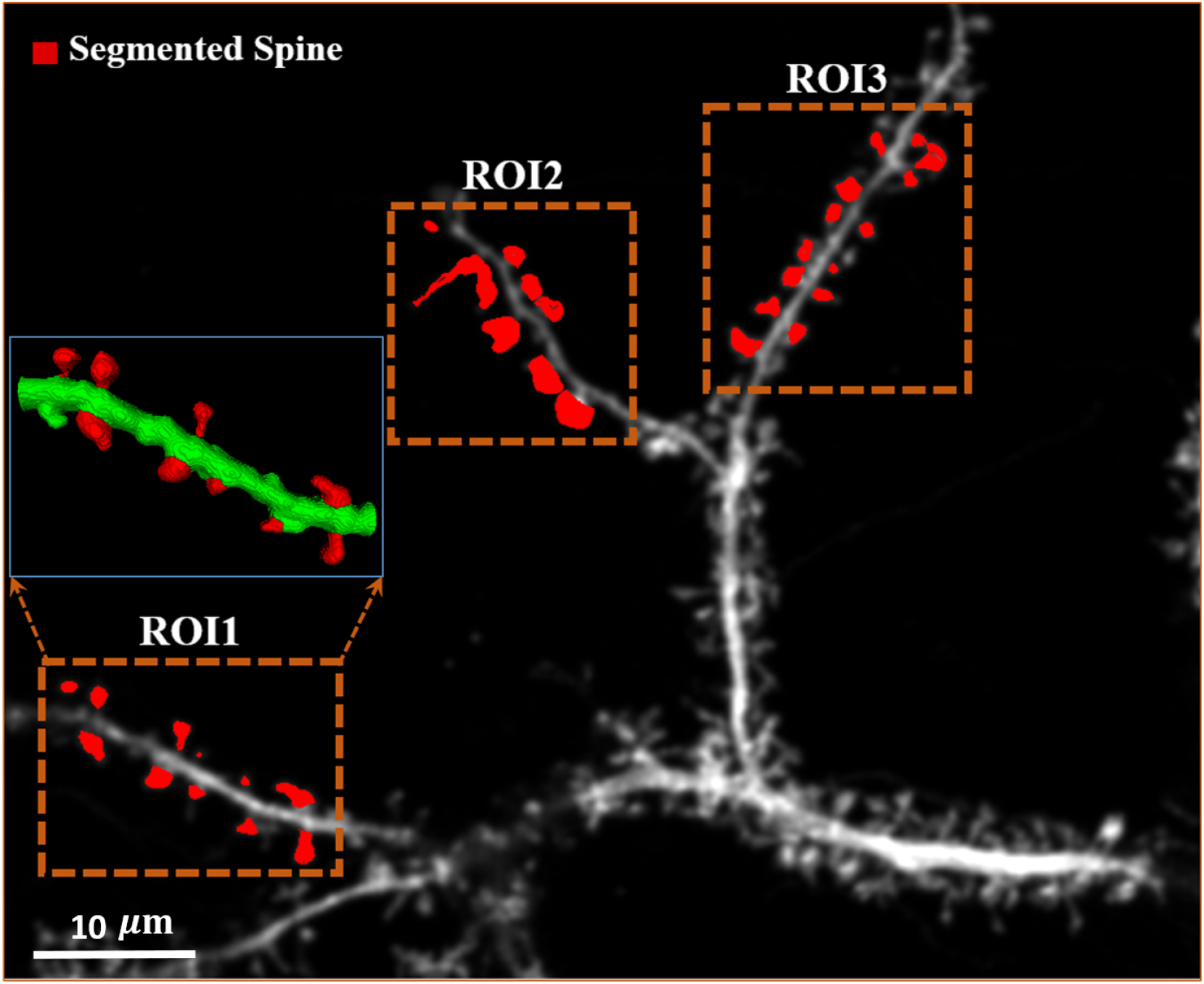
Different regions (ROI1, ROI2 and ROI3 are enclosed in dotted rectangular box) with segmented spines, confocal microscopic image of *in vitro* neuronal culture (Scale bar=10*μ*m). 3D rendering of the ROI 1 is shown in the inset. Segmented spines are shown in red color.

## 4. Conclusions

The main advantage of 3dSpAn software is its ability to segment and reconstruct individual spine in 3D for images different modalities and significant artifact contents, obtained using different laboratory techniques. The software extracts quantitative morphological features for dendritic spines and classifies them into one of the three classes: stubby, filopodia and mushroom spines. Each spine shape reflects the function and the strength of synaptic connections, thus precise determination of spine morphology is crucial in studies of synaptic plasticity [18]. The segmentation results were shown in *in vitro* and *ex vivo* images captured using confocal microscopy and *in vivo* image captured using two photon microscopy. The high reproducibility of proposed segmentation method has been already established in our previous publications [1,24].

## 5. Patents

Not Applicable

## Supplementary Materials

3dSpAn source code, 3dSpAn executable (3dSpAn V1.2 Installer for Windows), Manual(*3dSpAn_ Supplementary.pdf*), video tutorials, sample data are available online at: https://sites.google.com/view/3dSpAn/. The figures, tables etc. given in *3dSpAn_ Supplementary.pdf* are listed below.

**Figure S1:** The 3dSpAn GUI; **Figure S2:** A 3D image is loaded in the 3dSpAn software; **Figure S3:** Default Preprocessing option can be found in Tool menu of 3dSpAn GUI. Preprocessing settings dialog is shown in inset; **Figure S4:** Option to show Gridlines can be found in Display menu of the 3dSpAn GUI, **Figure S5:** Gridlines are displayed on the image loaded in 3dSpAn GUI; **Figure S6:**Options to select a region and crop that are enclosed with red boxes; **Figure S7:** Cropped region of the loaded 3D image is shown in the 3dSpAn GUI. The cropped region is named as ROI_1; **Figure S8:** The cropped ROI is magnified. The spin boxes enclosed by red box is used to tune the Pure Spine and Pure Dendrite intensity range; **Figure S9:** Option to show display the ROI as color-coded fuzzy segment can be found Display menu of the 3dSpAn GUI; **Figure S10:** The image is shown as color coded fuzzy segment depending on the pure spine and pure dendrite range; **Figure S11:** The image is shown as color coded fuzzy segment depending on the pure spine and pure dendrite range and the option to select seed type and add seed is enclosed by red boxes; **Figure S12:** A spine (encircled by red circle) is shown with pixel having intensity (119) greater than the shared intensity range upper bound (70) and falls in pure dendrite region; **Figure S13:** The upper bound of the shared intensity space between spine and dendrite is increased to 125.To see the added seeds, uncheck the “Fuzzy Segment” from Display–>Fuzzy Segment; **Figure S14:** The option to add spine/dendrite/separator can be found from Display–>Show Seed/Sep. Spine seeds are shown in red and dendrite seeds are shown in green; **Figure S15:** The option to Show Seed Id can be found from Display–>Show Seed Id–>Spine/Dendrite/Separator; **Figure S16:** To run the segmentation algorithm with user given seeds and separators user need to click on the Execute button, enclosed by red box; **Figure S17:** Segmented spines are shown in red and dendrite is shown in green; **Figure S18:** Click on Extract Feature (enclosed by red box) from the Analyse tab to calculated the morphological features of the segmented spine. After clicking Extract Feature segmented spine will be numbered in the order first spine seed marking; **Figure S19:** Check the option “Show Original Image” to go back to the full 3D image. Segmented spines are shown in red and segmented dendrite is shown in green; **Figure S20:** User can select a new ROI from the same image by checking the option Select a region and crop it by clicking on Crop button; **Figure S21:** The new ROI (ROI_2) is also segmented and numbered in the same way as ROI_1; **Figure S22**: The full image is shown with two segmented ROIs (ROI_1 and ROI_2). User can delete a ROI by putting ROI no. in the ROI No. and clicking Delete ROI button; **Figure S23:** After loading the image, if we load seeds/separators and previously cropped regions information from the option File–>Load All ROI Profile, then the segmented regions will shown enclosed by blue boxes; **Figure S24:** The ITK Snap GUI. New image can be loaded from File–>Open Main Image; **Figure S25**: The ITK Snap GUI. If we click File–>Open Main Image, then the dialog box will appear; **Figure S26:** The ITK Snap GUI. After selecting the image, choose Raw Binary from the File Format drop down; **Figure S27:** After selecting the image, input proper Image dimensions, Voxel type and click Next> button; **Figure S28:** Click Finish to complete the image loading; **Figure S29:** The loaded 3D image is shown in three different plane; **Figure S30:** To load the same image as segmentation go to Segmentation–>Open Segmentation; **Figure S31:** After loading the image as segmentation, spine and dendrite will be shown in different color (may not be always red and green). User can select the color for spine and dendrite intensity from color picker (enclosed by red circle). To see the 3D rendering click update button, enclosed by red box; **Figure S32:** All the three different plane views and 3D rendering window is shown together. To maximize 3D rendering window click on the 3D button enclosed by red box; **Figure S33:** 3D rendering of the segmented spine and dendrite.

**Table S1:** Volume, Length, Head Width and type of the segmented spines ROI_2 shown in the Figure S21.

**Video S1:** 3dSpAn Tuttorial-1; **Video S2:** 3dSpAn Tutorial-2; **Video S3:** 3D rendering of the segmented spine and dendrite.

## Author Contributions

S.B. and J.W. conceived the study; S.B., P.K.S., D.P., J.W. performed the experimental design; N.D., S.B. developed the software; N.D., E.B., M.B., A.Z., B.R. analyzed the data; S.B., N.D., B.R, E.B., J.W, E.P.,M.B. wrote the manuscript.

## Acknowledgments

Authors acknowledge CMATER Laboratory, Department of Computer Science and Engineering, Jadavpur University, Kol-32 for supporting the development.

## Abbreviations

MSO: Multiscale Opening
FDT: Fuzzy Distnace Transform
ROI: Region of Interest

## Appendix A

## Sample Availability

NA

© 2020 by the authors. Submitted to *Journal Not Specified* for possible open access publication under the terms and conditions of the Creative Commons Attribution (CC BY) license (http://creativecommons.org/licenses/by/4.0/).

